# The Haemagglutinin Genes of the UK Clade 2.3.4.4b H5N1 Avian Influenza Viruses from 2020 to 2022 Retain Strong Avian Phenotype

**DOI:** 10.1101/2024.07.09.602706

**Authors:** Jiayun Yang, Rebecca Daines, Pengxiang Chang, Thusitha K. Karunarathna, Mehnaz Qureshi, Jean-Remy Sadeyen, Joe James, Ashley C. Banyard, Marek Slomka, Ian H. Brown, Munir Iqbal

## Abstract

Since 2020, the United Kingdom (UK) has suffered repeated epizootics of clade 2.3.4.4b H5 high pathogenicity avian influenza viruses (HPAIVs) in wild birds and poultry, resulting in substantial economic losses due to enforced statutory control. The rapid evolution of H5 HPAIVs continues to raise concern with heightened zoonotic and pandemic risks. The immunodominant haemagglutinin glycoprotein (HA) is crucial for influenza virus receptor binding and pH-induced fusion of viral and cellular membranes. Mutations in HA are frequent due to polymerase error, immune pressure and host adaptation, resulting in antigenic modulation and/or an expansion of host tropism, respectively, ultimately hindering control strategies. We evaluated a comprehensive panel of H5 viruses representing prevalent genotypes from UK outbreaks spanning 2020 to 2022 for HA functionality. HA genes from each genotype were assessed through receptor binding, pH of fusion, thermostability and HA inhibition assays to evaluate factors contributing to zoonotic potential, stability, and antigenicity. The viruses only bound to avian receptors and exhibited fusion at a pH of 5.8, above the pH range (pH 5.0 to 5.5) associated with efficient human-to-human transmission. Therefore, these H5 viruses have low immediate zoonotic threat. Contemporary H5 viruses were more thermostable and showed antigenic drift compared to the earlier 2017-2018 clade 2.3.4.4b H5N8 viruses, and N236D in HA was identified as a significant antigenic epitope. The findings of this study underscore the evolving nature of the HA of these viruses and highlight the importance of ongoing surveillance and characterisation efforts to identify factors that might contribute to zoonotic risk.

## Introduction

Influenza A virus (IAV) is a negative-sense, single stranded RNA virus with a genome consisting of eight gene segments. The natural reservoirs for influenza A viruses are aquatic waterfowl, namely Anseriformes, where they are referred to as avian influenza viruses (AIVs), although some subtypes can infect a wide range of avian and mammalian hosts, including humans. AIV subtypes are classified based on the antigenicity of the viral glycoproteins haemagglutinin (HA) and neuraminidase (NA), with 16 HA subtypes and 9 NA subtypes widely identified [1, 2]. Since the emergence of the A/goose/Guangdong/1/96 (Gs/GD/96) lineage of H5 high pathogenicity avian influenza virus (HPAIV), the global poultry industry has continuously been economically devastated, driven by the migratory behaviour of wild bird populations. The Gs/GD/96-like viruses have also raised concerns for public health, with 461 human deaths from 882 infections (52.3%) being reported since 2003 [3, 4]. To date, despite a general decrease in zoonotic threat due to continual genetic reassortment, human infections with HPAIV H5 viruses remain sporadically reported [5].

The HA surface glycoprotein initiates viral infection in host cells and is also immunodominant and thus modulates antigenicity although the other key glycoprotein, the neuraminidase protein (NA) also plays a role in antigenicity. Since its emergence, the HA of the Gs/GD/96 lineage has evolved into over 30 clades and subclades resulting in substantial antigenic diversity [6]. From these clades, clade 2.3.4.4 has become dominant in recent years and has evolved antigenically into eight additional subclades, 2.3.4.4a to h [7]. Since its emergence by 2016, the dominant subclade 2.3.4.4b has become disseminated globally since 2019 [8–10], with multiple genotypes emerging, termed H5Nx (where Nx is N1-N9), with various phenotypic adaptations in domestic and wild birds [11]. The first of the major clade 2.3.4.4b epizootics detected in the United Kingdom (UK) was on 2^nd^ November 2020, and since then the UK has continued to see numerous infections in domestic poultry and wild bird populations [12][13]. In addition to the unprecedented avian outbreaks caused by the contemporary clade 2.3.4.4b H5 viruses, numerous and increasing numbers of mammalian cases are being reported globally including seals, minks, cats, and recently dairy cows [14–17]. Most of these infections are thought to the result of exposure via environmental contamination or predation upon infected birds [18]. Furthermore, sporadic human infections caused by 2.3.4.4b have been reported, typically the result of close contact with infected poultry [19]. To date, mammal to mammal transmission has been suspected although conclusive evidence demonstrating this hasn’t been reported. Further, from a human perspective, despite human detections no human-to-human transmission has been reported.

Functionally, the viral HA recognises and binds to the sialic acid receptors on the cell membrane to initiate infection of host cells through endocytosis, facilitating entry through fusion of the cell and viral membranes [20]. The binding of HA to sialic acids drives host tropism, where avian adaptation is reflected by a preferential affinity of the HA of avian viruses to α-2,3-sialic acid (α-2,3-SA), while human or mammalian adaptation is reflected typically by a high affinity of binding to α-2,6-sialic acid (α-2,6-SA) [21–23]. Typically, Has of human viruses require lower pH (pH<5.5) for subsequent endosomal membrane fusion, whereas avian viruses require pH>5.5 [24].

Despite an unprecedented expansion of both avian and mammalian infections with contemporary H5N1 clade 2.3.4.4b viruses there remains limited knowledge on mammalian adaptive mutations that may increase zoonotic threat. Therefore, we investigated the functional capacity of the HAs of these viruses through receptor binding profiles, thermal and pH stability, and antigenic variation, comparing contemporary viruses with pre-2020 predecessor clade 2.3.4.4b viruses.

## Materials and Methods

### Ethics statement

All the procedures involving embryonated eggs were undertaken in strict accordance with the guidance and regulations of the UK Home Office under project licence number PP6471846. As part of this process, the work has undergone scrutiny and approval by the animal welfare ethical review board at The Pirbright Institute, incorporating the 3Rs and followed by ARRIVE (Animal Research: Reporting of *in vivo* experiment) guidelines for quality, reproducibility and translatability of animal studies.

### Cells and viruses

Madin-Darby canine kidney (MDCK) cells, human embryonic kidney 293 (HEK-293) cells, and African green monkey kidney epithelial (Vero) cells were maintained by and acquired from the Central Services Unit (CSU) of TPI. Cells were maintained with Dulbecco’s Modified Eagle’s medium (DMEM) supplemented with 10% fetal calf serum (FCS) (Gibco) and maintained at 37°C with 5% CO_2_.

The HA sequences of two predecessor H5N8 viruses A/crow/Aghakhan/2017 (EPI_ISL_320957) and A/chicken/Cheboksary/854/2018 (EPI_ISL_380991) were acquired from the Global Initiative on Sharing All Influenza Data (GISAID). Additionally, the HA and NA sequences from the recent dominant genotypes of H5Nx viruses isolated within the UK were provided by the Animal and Plant Health Agency (AHPA). Viruses include: A/chicken/England/030786/2020 (H5N8) (EPI_ISL_17212363), A/mute swan/England/234255/2020 (H5N1) (EPI_ISL_766876), A/chicken/England/053052/2021 (H5N1) (EPI_ISL_9012457), A/chicken/Wales/053969/2021 (H5N1) (EPI_ISL_9012618), A/chicken/Scotland/054477/2021 (H5N1) (EPI_ISL_9012696), A/chicken/England/085598/2022 (H5N8). Abbreviations of the H5Nx viruses used were named as Agha/17, Che854/18, Eng786/20, Eng255/20, Eng052/21, Wal969/21, Sct466/21 and Eng598/22, respectively. The polybasic cleavage site of H5 HAs PLRERRRKR/GLF was replaced with a monobasic PLGTR/GLF cleavage site for handling at containment level 2 (CL2). Each HA and NA sequence were synthesised by GenScript and cloned into a pHW2000 plasmid vector. Viruses were rescued by reverse genetics (RG) using an eight-plasmid bidirectional expression system [25] with the internal segments from laboratory adapted strain A/Puerto Rico/8/1934 (H1N1) (PR8). The RG viruses were propagated in 9 to 10-day old embryonated hen’s eggs for working and purification stocks. A reassortant H9N9 virus containing HA of A/Chicken/Pakistan/UDL-01/2008 (H9N2) and NA of A/Anhui/01/13 (H7N9) was used as a control virus for receptor binding analysis [26]. The Eng052/21 HA plasmid with D236N mutation was generated using the single-site mutagenesis QuikChange II (Agilent) kit following manufacturer’s instructions, and primers (5’ to 3’) forward: CTCTCGAAATGGATTGCATCATTTGGTTTTAAAATTGTCCAGAAGA and reverse: TCTTCTGGACAATTTTAAAACCAAATGATGCAATCCATTTCGAGAG.

### Virus purification and quantification

Propagated viruses were purified as described previously [27]. In brief, 400mL of virus in allantoic fluid were concentrated by ultracentrifugation at 27,000 rpm for 2 hours at 4[. Virus pellets were homogenised and purified through a 30% to 60% sucrose gradient at 27,000 rpm for 2 hours at 4[, with a reduced rate of deceleration. Purified virus was again pelleted to remove sucrose at 27,000 rpm for 2 hours at 4 [and reconstituted in phosphate buffered saline (PBS). Enzyme-linked immunosorbent assay (ELISA) was used for purified virus quantification as described previously [28] by comparing viral nucleoprotein (NP) detection to a known standard.

### Bio-layer interferometry (BLI)

Avian-like sugar analogues α-2,3-sialyllactosamine (3SLN) and Neu5Ac α-2,3Galβ1-4(6-HSO3) GlcNAc (3SLN (6su)) and human-like sugar analogue α2,6-sialyllactosamine (6SLN) were purchased from GlycoNZ. Receptor binding was assessed with BLI methods as previously described [29]. In brief, 100 pmol of purified virus was diluted with HBS-EP buffer (TEKnova) with additional 10 μM oseltamivir carboxylate (Roche) and 10 μM zanamivir (GSK) to exclude sialidase activity of NA; the receptor binding against each sugar analogue was tested with Octet® R8 system (Sartorius) using streptavidin biosensors (Sartorius). Virus binding affinity was normalised to fractional saturation and the concentrations of sugar loadings [30].

### Haemagglutination assay and haemagglutinin inhibition (HI) assay

Haemagglutination assay and haemagglutinin inhibition (HI) assay were performed following World Health Organisation (WHO) guidelines, using 1% (0.5% final volume) chicken red blood cells (RBCs) [31]. Heat-inactivated post-infection duck serum, collected from ducks infected with Eng052/21 at a dilution of 1:4 (v/v), were used in the HI assay [32]. All viral titres were calculated to haemagglutinating units (HAU)/50 µL.

### Thermostability assay

The thermostability assay was tested as previously described [33]. HA titres were diluted with allantoic fluid and normalised to 32 HAU/50 µL. The viruses were then incubated in the Thermal Cycler (Bio-Rad) at 50°C, 50.7°C, 51.9°C, 53.8°C, 56.1°C, 58.0°C, 59.2°C, and 60°C and 4°C as control for 30 minutes. The thermostability of the incubated viruses were then determined by haemagglutination assay.

### Syncytium formation assays

Syncytium formation assays were performed as previously described [33]. In brief, viruses were titrated in Vero cells by using anti-nucleoprotein (anti-NP) mouse monoclonal antibody and horseradish peroxidase-labelled rabbit anti-mouse immunoglobulins (Dako). Monolayered Vero cells were then infected with virus at a multiplicity of infection (MOI) of 1 for 16 hours. The cells were then treated with 3.0 μg/mL N-tosyl-L-phenylalanine chloromethyl ketone (TPCK) trypsin for 15 minutes, followed by incubation with PBS ranging from pH 5.2 to 6.0 at 0.1 pH increment for 5 minutes. The cells were maintained with DMEM+10%FCS for 3 hours and fixed with acetone/methanol (1:1 ratio) and stained with Giemsa stain (Sigma-Aldrich). Stained cell images were taken by EVOS XL imaging system (Life Technologies).

### Phylogenetic analysis

Sequences were sourced (WHO/WOAH/FAO, 2014) for clades 1 – 9 except 2.3.4.4 and the 2020 CDC risk assessment for clade 2.3.4.4 viruses [34]. The phylogenetic tree was initially constructed as neighbour-joining to predict the most accurate tree model for maximum-likelihood (ML) by using MEGA11 [35, 36]. The ML tree was then constructed using the GTR+G substitution model selected for the lowest Bayesian Information Criterion (BIC) from the best-fit model, assuming equal rates of evolution, a fixed rate of heterogeneity across sites and rooted at the mid-point. The classification and assignment of clades of H5Nx viruses assumes a common ancestral node and monophyletic evolution with a bootstrap value ≥ 60 at the defining nodes, following 1000 bootstrap replicates [37]. The tree was annotated in R Studio (R version 4.3.2) using the ggtree package (version 3.6.2) [38].

### Haemagglutinin structure prediction

The crystal structure of H5 HA was acquired from Protein Data Bank (https://www.rcsb.org/) with assession number 4JUL. HA structure modelling and prediction were visualised using SWISS-MODEL and visualised and annotated in PyMol version 4.6.

### Statistical analysis

Data were analysed and visualised by GraphPad Prism 10.0 (GraphPad Software, USA). Significance was determined by two-way ANOVA multiple comparisons and Ordinary one-way ANOVA Dunnett’s multiple comparisons test. Levels of significance (p) are denoted as, 0.01<p<0.05 having one asterisk, and 0.001<p<0.01, 0.0001<p<0.001 and p<0.0001 having two, three or four asterisks, respectively. p>0.05 was considered not significant.

## Results

### Molecular characterisation of HA of H5Nx

A panel of six viruses, including one H5N8 virus, Eng786/20, and five H5N1 viruses, Eng255/20, Eng052/21, Wal969/21, Sct477/21 and Eng598/22 were selected based on the epizootic timelines of the dominant genotypes detected in the UK from 2020 to 2022 [39]. Eng786/20 and Eng255/20 were identified in 2020 as the precursor H5N8 and H5N1 viruses, soon both viruses were replaced by Eng052/21 and Wal969/21 in 2021, and then Sct477/21 became the most dominant genotype until 2022 when Eng598/22 emerged [39]. Phylogenetic analysis of the HA gene confirmed that the selected H5Nx viruses belong to clade 2.3.4.4b (Figure 1A). All the HA numbering system used is based on H5 numbering. Details of the amino acid variations within the HA of the study viruses are presented in Figure 1B. The HA molecular markers that have previously shown increased binding to α-2,6-SA and increased virulence in mammalian hosts of H5 viruses (at amino acid (aa) position 133, 188, 196, 197, 222 and 226) are included in *Table S1*. The six study H5Nx viruses have Alanine (A), Threonine (T), Lysine (K), Asparagine (N), Glutamine (Q) and Glycine (G) at aa position 133, 188, 196, 197, 222 and 226, respectively (H5 numbering used henceforth). HA aa mutations of S133A, T188I, Q196R/H, N197K, Q222L and G226S are related with increased binding affinity to human-type receptors than avian-type receptors in H5 viruses [40–43]. Apart from aa 133, the other five aa positions showed binding affinity to avian-type receptor. All the H5Nx viruses used in this study show H at aa 103 and A at aa 156, mutations of H103Y and T156A were shown to have increased H5N1 transmissibility in ferrets [40]. Three HA mutations (I151T, G154N and W221G) that are associated with mammalian adaptations including increased binding to human lower respiratory tract, increased replication in mammalian cells and increased virulence in mice, are also present in the selected H5Nx viruses [43–45]. In summary, the HAs of all chosen viruses have conservation across the key amino acids, and this observation implied that the HA of current clade 2.3.4.4b H5 viruses have acquired a few amino acid mutations that may be related to mammalian adaptation.

**Figure 1.**
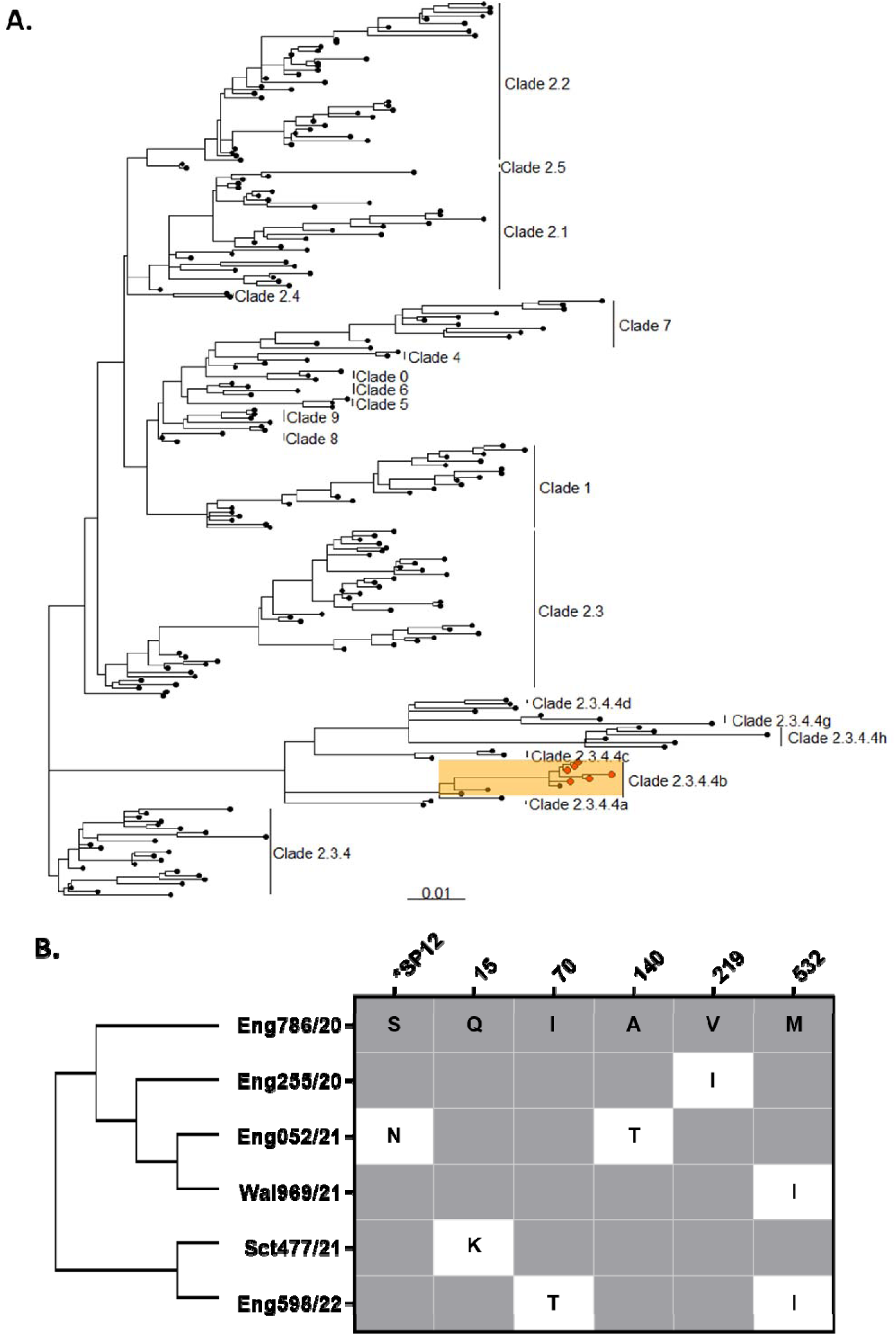
Phylogenetic analysis and amino acid divergence of HA from studied viruses. (A) Viral HA maximum-likelihood (ML) tree constructed using GTR-G substitution model. The classification and assignment of H5 clades were based on bootstrap value ≥ 60 at defining nodes, following 1000 bootstrap replicates. The viruses used in this study (red) belong to clade 2.3.4.4b (orange highlight). (B) HA amino acid divergence of study viruses (by H5 numbering) with conserved amino acids (grey), and divergent amino acids (white). *SP indicates signalling peptide of HA.

### H5Nx virus isolates from the UK have a high affinity to avian receptors

The binding profiles of the H5Nx viruses against human and avian receptor analogues were assessed by BLI. Simple sialylated avian-like analogue 3SLN and human-like receptor analogue 6SLN were used to assess the receptor binding profiles of the H5Nx viruses. In addition, 3SLN(6Su) was also added for receptor binding assessment as our previous data showed that western G1 lineage H9N2 viruses bound to 3SLN(6Su) but not 3SLN [29]. The reassortant H9N9 virus that showed binding to 3SLN, 3SLN(6Su) and 6SLN was used as a positive control (Figure 2A) [26]. None of the H5Nx viruses bound to human-like receptor analogue 6SLN, but showed clear binding to avian receptor analogues 3SLN and 3SLN(6Su) (Figure 2B to 2G), with the amino acid differences and dissociation constant (*K_d_*) values for virus receptor binding listed in *Table S2*. Eng052/21, Eng255/20, Sct477/21, Wal969/21, and Eng598/22 showed weaker binding to 3SLN (Figure 2H) and 3SLN(6Su) (Figure 2I) in comparison to Eng786/20.

**Figure 2.**
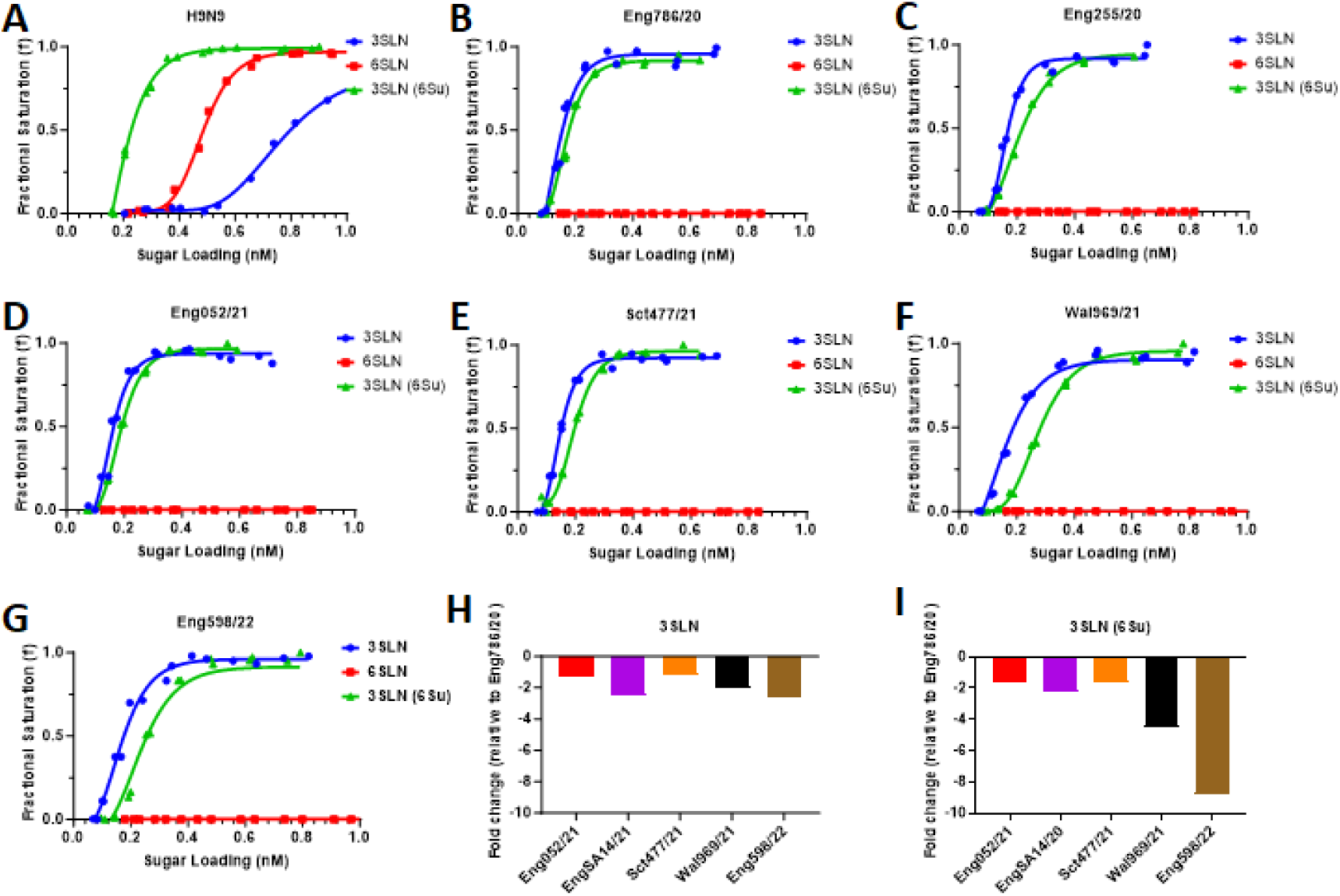
Receptor-binding profiles of the clade 2.3.4.4b H5Nx viruses. BLI was used to assess virus receptor binding to avian-like receptor analogues (3SLN and 3SLN (6Su)) and human-like receptor analogue (6SLN) on the following viruses: (A) H9N9 as positive assay control, which shows binding to all three receptor analogues, (B) Eng786/20, (C) Eng255/20, (D) Eng052/21, (E) Sct477/21, (F) Wal969/21, and (G) Eng598/22. The fold change of 3SLN and 3SLN(6Su) relative to Eng786/20 are indicated in (H) and (I), respectively.

### The UK H5Nx viruses show membrane fusion at pH 5.8

HA undergoes a conformational change at low pH into its active form that results in endosomal membrane fusion during the infection process [46]. The fusion pH range from 5.0 to 5.9 is associated with AIVs which are able to establish efficient infection and transmission among birds, but for mammalian species, in particular humans, the HA requires a lower pH for fusion (pH5.0 to 5.5) [47, 48]. Vero E6 cells were infected with the H5 viruses and then treated with TPCK trypsin to instigate fusion. All six reverse genetics generated H5 viruses, Eng786/20, Eng052/21, Eng255/20, Sct477/21, Wal969/21, and Eng598/22, demonstrated syncytium formation at a pH of 5.8 (Figure 3).

**Figure 3.**
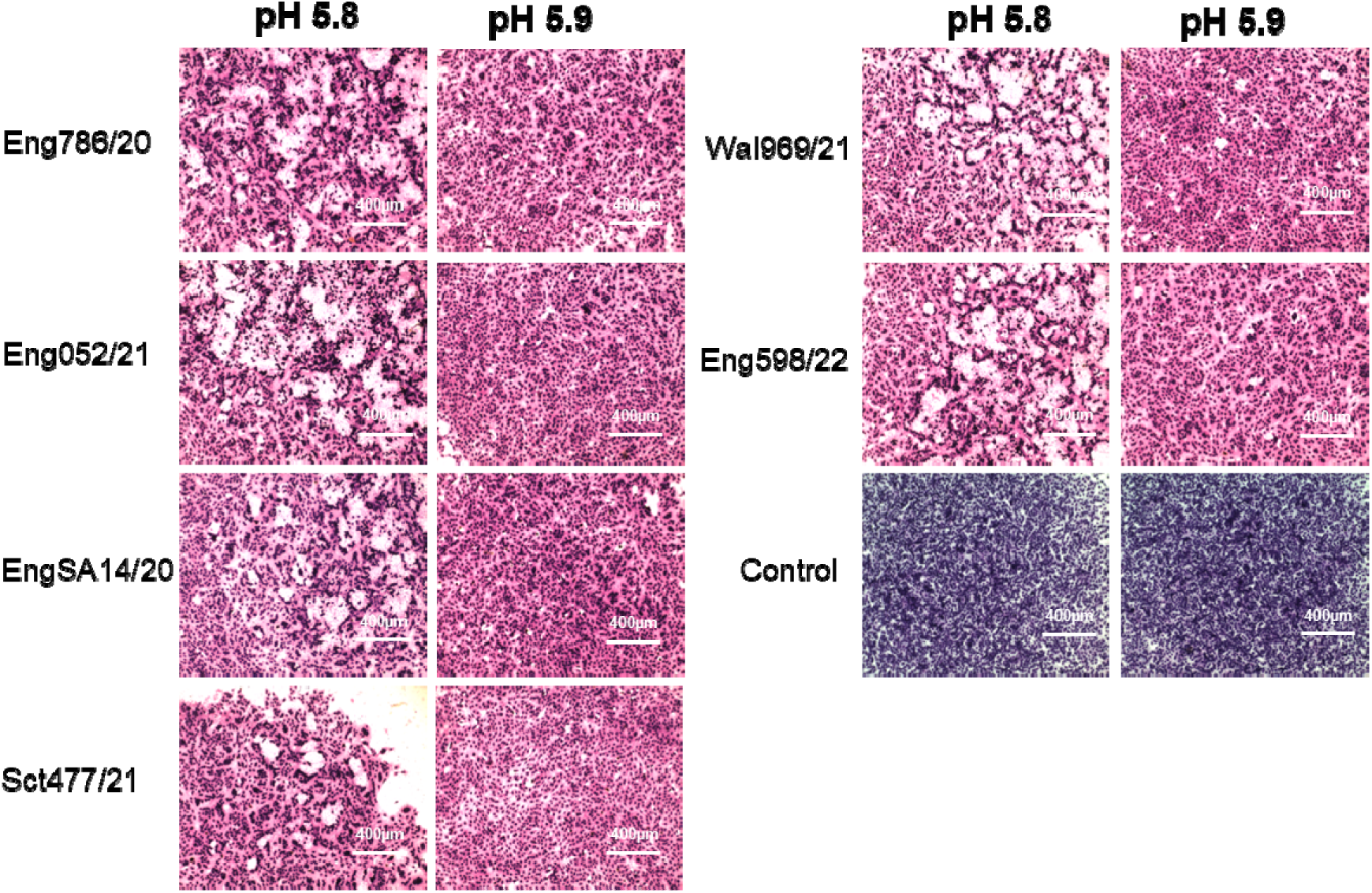
pH of fusion of clade 2.3.4.4b H5Nx viruses from the UK. Syncytium formation assays were used to evaluate initiation of fusion of panel viruses by infecting monolayered Vero cells with a range of pH treatments. Fusion pH was determined based on 50% of maximum syncytium formation. The images were captured at scale of 400 µm.

### Contemporary UK H5 viruses show increased thermostability compared to previous H5N8 viruses

The thermostability of the H5 viruses was assessed to test HA stability. Two earlier H5N8 viruses of clade 2.3.4.4b H5N8, Agha/17 and Che854/18 were added for comparison with UK H5Nx viruses. Viruses were diluted to 64 HAU and heat-treated at a range of temperatures from 50°C to 60°C, with 4°C serving as a control. In comparison to previous H5N8 virus Che854/18, the UK clade 2.3.4.4b viruses were more stable at higher temperatures, since Che854/18 lost HA activity completely at 50°C (Figure 4). The thermostability of Agha/17 reduced 4-fold at temperatures at 51.9°C, whereas most of the current H5 viruses remained unchanged. Among the six current H5 viruses, Eng598/22 showed a more significant HAU reduction than other viruses at between 51.9 and 53.8°C (p<0.0001).

**Figure 4.**
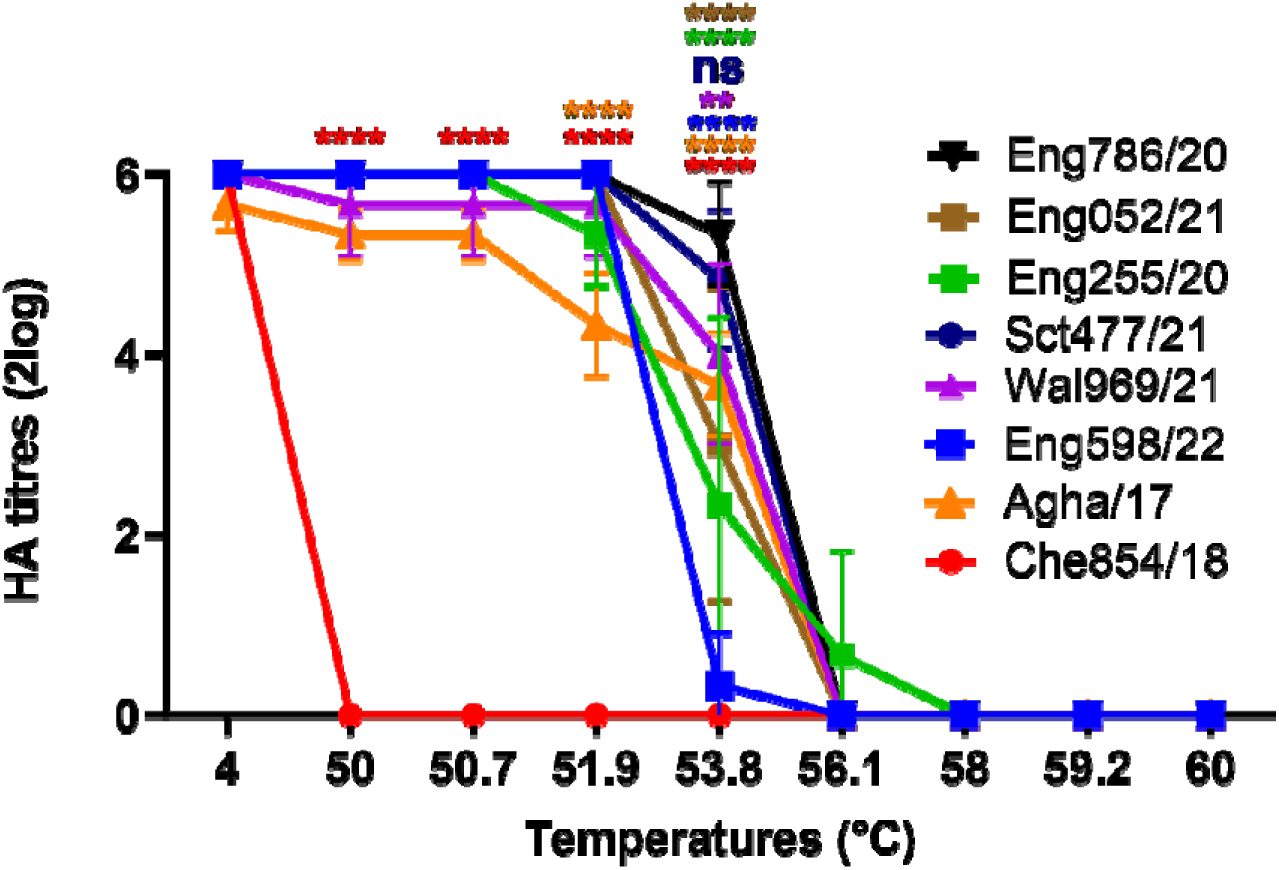
Thermostability of 2020-2022 UK clade 2.3.4.4b H5Nx viruses in comparison to 2017-2018 H5N8 viruses. Input virus was standardised to 64 HAU and heat-treated at the temperatures indicated above for 30 minutes; 4°C served as a negative control. HA assays were then performed after heat treatment to observe HA stability through erythrocyte agglutination. The experiment was repeated three times independently, and significance indicated on those with reduction in HA titres in comparison to the other viruses, ns = not significant, ** = p<0.01, **** = p<0.0001.

### The N236D mutation was identified as epitope that altered antigenicity

Homologous post-infection duck antiserum of Eng052/21 were used in the HI assay to assess antigenicity (Figure 5A). The five heterologous H5 viruses (Eng786/20, Eng255/20, Sct477/21, Wal969/21, and Eng598/22) had comparable HI titres (∼7log2) in comparison to homologous Eng052/21 virus. Coincidentally, the HA of the WHO candidate vaccine virus (CVV) A/Astrakhan/3212/2020 (H5N8) shared identical amino acid sequences with Eng786/20. Two previous clade 2.3.4.4b viruses, Che854/18 and Agha/17, the mean values of HI titres showed 2 and 1 log2 reduction, respectively. One mutation at amino acid position 236 from Aspartic acid (D) to Asparagine N (D236N) located within the antigenic globular head region (Figure 5B; *Table S3*), which could be the potential epitope driving antigenicity [42, 49]. The mutation was introduced to the Eng052/21 RG virus and a resultant antigenic change observed by the HI assay. The HI titre of D236N mutant virus was lower than WT Eng052/21 virus by 1 to 3 log2-fold in three biological repeats (Figure 5C).

**Figure 5.**
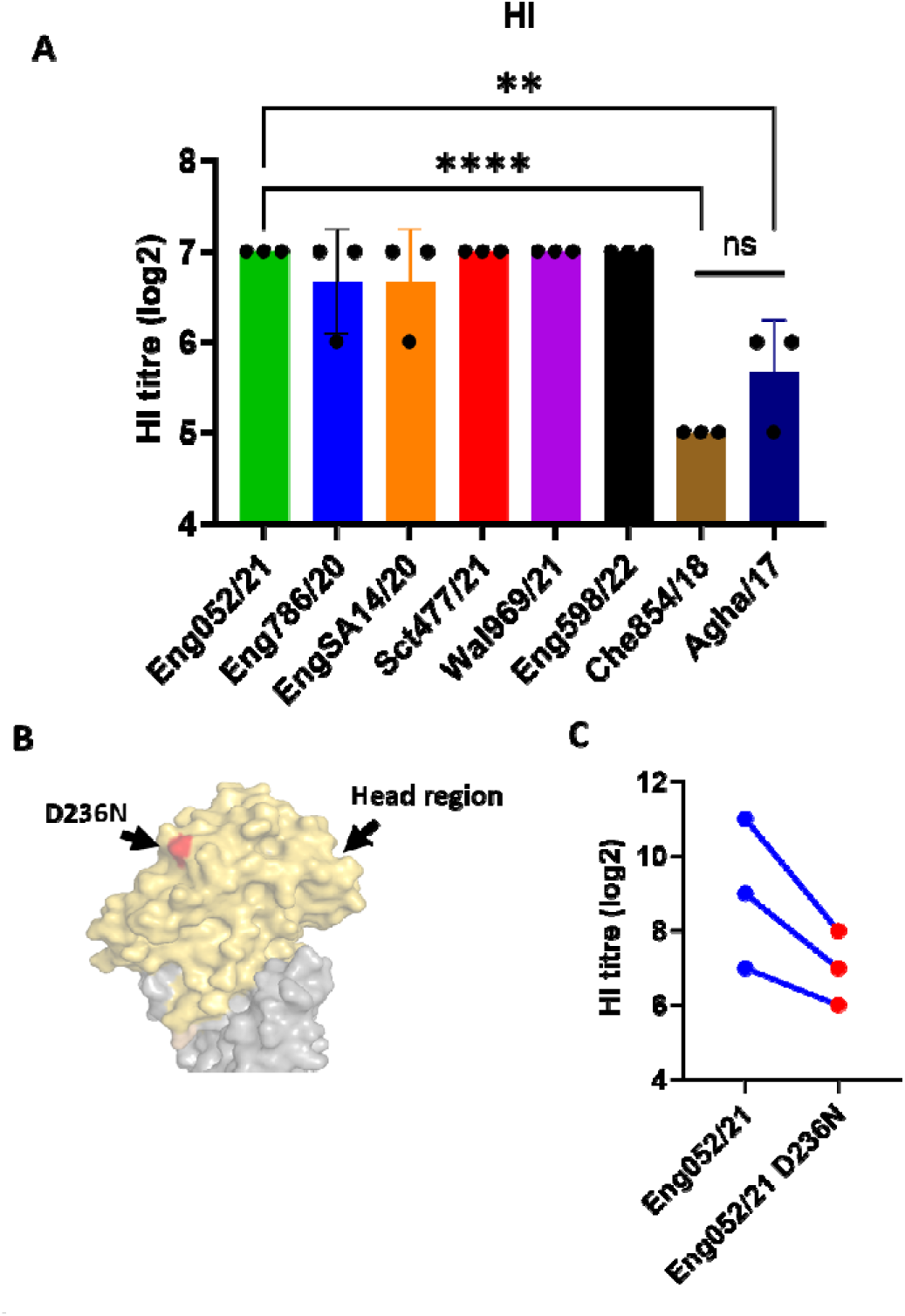
Antigenicity assessment between contemporary 2020-2022 and previous 2017-2018 H5Nx viruses. (A) 2017-2018 H5N8 viruses show decreased HI titres compared with contemporary H5Nx viruses. Levels of significance are shown in figures, ** = 0.001<p<0.01, **** = p<0.0001, ns = not significant. (B) N236D (red) location within the head region (yellow) of HA1. HA structure was generated using template HA of clade 2.3.4 A/duck/Laos/3295/2006 (H5N1) (PDB: 4JUL). (C) Eng052/21 with D236N mutation shows decreased HI titre compared with Eng052/21. HI assays were performed using duck antisera raised against Eng052/21 with three biological repeats.

## Discussion

The contemporary clade 2.3.4.4b H5 HPAIVs have become panzootic, and have been widly identified across all continents, including Antarctica during 2022-2023, surpassing the magnitude of all previous HPAIV outbreaks [50]. They pose a credible risk to wildlife and the poultry industry but with a potential, but currently undefined risk to public health. The clade 2.3.4.4b viruses have undergone serial reassortments and adaptations in bird populations with notable genotypic changes, in particular the shift of NA from N8 to N1 [51]. Further, the viruses causing the recent panzootic have undergo significant reassortment of the internal gene segments [39]. Influenza virus reassortments and adaptations pose great risk to human health. Recent pandemic and epidemic influenza A viruses, including the 2009 H1N1 pandemic [52] and the epizootic H7N9 virus in China [53], have featured reassortants with genetic segments with genetic segments of avian origin [54]. Therefore assessment of viral emergence and in particular evolution of HA is critical to define zoonotic risk.

The binding of viral HA to host cell sialic acid receptors is a prerequisite for AIV infection. Our data showed that the UK clade 2.3.4.4b H5 viruses have no detectable binding affinity to the human receptor, but showed strong yet moderately variable binding to avian receptors. Despite having four mutations (S133A, G154N, T156A, and W221G) considered relevant to mammalian adaptation [40, 42, 44, 45], none of the H5 viruses assessed showed increased binding affinity to human-like receptor 6SLN, with all retaining highest binding affinity to the avian-like receptor analogues 3SLN and 3SLN(6Su).

Sulphated Neu5Acα2-3Gal was discovered widely distributed in terrestrial birds including chickens and quails, but were absent from ducks, indicating sulphation of sialic acid appears to play a key role of host range in avian species [55]. Interestingly, the recent global outbreaks of H5N1 have manifest on numerous occasions in mammalian spillover infections [14, 16]. Whilst these are of concern, the receptor binding data presented here suggests that the HA circulating in the UK from 2020 to 2022 posed limited zoonotic risk, and this aligns with global sequence obtained from mammalian species where very limited adaptation in HA has been noted. However, although there has been a lack of HA adaptation, several H5N1 isolates from wild mammals, including terrestrial and marine mammals, have acquired mammalian adaptations such as PB2 Q591K and M535I [56, 57]. Further, a recent study has shown a highly pathogenic North American strain (Scaup/GA/22) was able to transmit among ferrets [58]. As such, the surveillance and assessment of currently circulating H5 viruses is a high priority for pre-pandemic preparedness.

The acid stability of the viral HA protein also plays a crucial role in virus transmission and the host range for efficient virus transmission appears to be pH dependent. In this study, we tested the fusion of clade 2.3.4.4b H5Nx viruses, and our data demonstrated syncytium formation at a pH of 5.8. This is consistent with a recent study that demonstrated that North America clade 2.3.4.4b H5N1 viruses had a pH of fusion at 5.8 [58]. This correlates with the fact that human influenza viruses require lower pH for membrane fusion pH≤5.5. For example, pandemic viruses such as H1N1 (2009) and H3N2 (1968) viruses were active at pH 5.46 and 5.25 respectively, whereas avian influenza viruses require higher pH of fusion (pH>5.5) [29]. The high pH of fusion (pH>5.5) showed that the current clade 2.3.4.4b H5Nx HPAIVs maintained this feature of avian-adapted viruses.

Thermostability has also been shown to be crucial for H5 AIV transmissibility in mammals. An HA mutation H103Y in the H5N1 virus contributed to the transmissibility in ferrets with increased HA thermostability and binding to human-like receptors [59]. In the current study, the HA of contemporary clade 2.3.4.4b H5Nx viruses has demonstrated higher thermostability than the previous H5N8 viruses at temperatures above 50°C. This showed that the current clade 2.3.4.4b viruses are more thermal stable than previous clade 2.3.4.4b H5N8 viruses. However, the consequences of whole viral thermostability which may influence survival in the environment may be a product of not only a more thermostable HA binding, but may include other contributory factors effected by temperature, such as viral polymerase activity and/or internal virion structure [60]. The extensive viral contamination of the environment by ducks experimentally infected by H5Nx clade 2.3.4.4b HPAIVs has been demonstrated [32, 61], indicating that virus persistence in the environment likely plays an important role in sustaining epizootic-scale infection pressure, stemming from infected wild waterfowl. These observations have been supported by detection of H5N1 clade 2.3.4.4b HPAIV in environmental specimens collected during UK poultry (including commercial ducks) outbreaks in 2023 [62].

It was hypothesised that HA mutation N236D in current clade 2.3.4.4b H5 virus is a possible epitope, as position 236 is in the highly variable and antigenic globular head of HA1 (Figure 5C) [63]. In this study, we compared the antigenicity between 2020-2022 clade 2.3.4.4b H5Nx viruses with 2017-2018 H5N8 viruses; the HI results confirmed that N236D mutation in the receptor binding site drove antigenic drift. Additionally, phylogenetic analysis indicated that the mutation emerged in domestic birds between 2017-2020 [11]. Therefore, N236D mutation is likely to have been an antigenic mutation between previous 2017-2018 clade 2.3.4.4b H5 viruses and contemporary 2020-2022 UK H5 viruses. Interestingly, Eng786/20 and Eng052/21 showed no antigenic difference (Figure 5) despite having two amino acids difference (Figure 1B), aa140 and aa219, which fall within the receptor binding site (RBS), are not antigenic epitopes.

In conclusion, our data suggests that the HA protein of 2020-2022 clade 2.3.4.4b H5Nx in the UK showed: i) preferential binding to avian-like receptor analogues; ii) and initiation of fusion at pH 5.8, which is a typical pH for fusion seen with avian viruses, ultimately suggesting the viruses pose limited risk to humans. Additionally, the contemporary H5Nx viruses were also more stable at higher temperature than 2017-2018 H5N8 viruses. Although the contemporary 2020-2022 viruses and previous 2017-2018 clade 2.3.4.4b viruses were genetically and antigenically similar, the HA mutation N236D within contemporary strains was identified to have contributed to antigenic drift in comparison to previous 2017-2018 H5N8 viruses. Ultimately, the surveillance and risk assessments of currently circulating strains remain crucial for understanding the potential threats posed by emerging viruses and for informing strategies to mitigate their impact on public health and agricultural sectors.

## Acknowledgements

We gratefully acknowledge all data contributors, i.e., the authors and their originating laboratories responsible for obtaining the specimens, and their submitting laboratories for generating the genetic sequence and metadata and sharing via the GISAID Initiative, on which this research is based. All submitters of the data may be contacted directly via the GISAID website (https://www.gisaid.org). We thank Dr. Steve Martin for sharing Octet analysis software for receptor binding analysis.

---

**Table S1.** HA molecular characteristics (by H5 numbering) of selected H5Nx viruses. “-” indicates the amino acid is the same as Eng786/20.

| Amino acid | Eng786/20 | Eng255/20 | Eng052/21 | Sct477/21 | Wal969/21 | Eng598/22 | Phenotype change |
| --- | --- | --- | --- | --- | --- | --- | --- |
| H103Y | H | - | - | - | - | - | Increased transmissibility in ferrets. |
| S133A | A | - | - | - | - | - | Increased recognition of α-2,6-SA by glycan array. |
| I151T | I | - | - | - | - | - | Increased binding to human lower respiratory tract. |
| G154N | N | - | - | - | - | - | Increased replication in mammalian cells. |
| T156A | A | - | - | - | - | - | Increased binding to α-2,3-SA and α-2,6-SA. |
| T188I | T | - | - | - | - | - | Increased recognition of α-2,6-SA by glycan array. |
| Q196R | K | - | - | - | - | - | Increased binding to α-2,6-SA. |
| Q196H | K | - | - | - | - | - | Increased binding to human lower respiratory tract. |
| N197K | N | - | - | - | - | - | Increased binding to α-2,6-SA. |
| W221G | G | - | - | - | - | - | Increased virulence in mice. |
| Q222L | Q | - | - | - | - | - | Increased binding to α-2,6-SA, Increased transmissibility in ferrets. |
| G226S | G | - | - | - | - | - | Increased binding to α-2,6-SA, Increased transmissibility in ferrets. |

**Table S2.** Dissociation constant values (*K_d_*) for virus receptor binding.

| Viruses | Absolute estimated KD values (nM) |  |
| --- | --- | --- |
|  | 3SLN | 3SLN (6su) |
| Eng786/20 | 113.7 | 328.3 |
| Eng255/20 | 283.9 | 733.8 |
| Eng052/21 | 156.7 | 535.5 |
| Wal969/21 | 235.6 | 2891.2 |
| Sct477/21 | 134.7 | 550.1 |
| Eng598/22 | 305.5 | 1493.5 |

**Table S3.** HA amino acid differences (by H5 numbering) between contemporary 2020-2022 UK H5Nx viruses and previous 2017-18 H5N8 viruses. Epitope N236D is highlighted (red). “-” indicates the amino acid is the same with Eng052/21.

| Viruses | Amino Acid Variance (H5 numbering) |  |  |  |  |  |  |  |  |  |  |  |  |  |  |  |  |  |
| --- | --- | --- | --- | --- | --- | --- | --- | --- | --- | --- | --- | --- | --- | --- | --- | --- | --- | --- |
|  | 15 | 71 | 82 | 97 | 138 | 140 | 219 | 236 | 268 | 277 | 320 | 322 | 376 | 378 | 491 | 523 | 528 | 533 |
| **Eng052/21** | Q | I | R | D | Q | T | V | **D** | G | K | S | L | D | V | Q | A | A | M |
| **Eng255/20** | - | T | - | - | - | A | - | - | - | - | - | - | - | - | - | - | - | I |
| **Eng786/20** | - | - | - | - | - | A | - | - | - | - | - | - | - | - | - | - | - | - |
| **Sct477/21** | K | - | - | - | - | A | - | - | - | - | - | - | - | - | - | - | - | I |
| **Wal969/21** | - | - | - | - | - | A | - | - | - | - | - | - | - | - | - | - | - | - |
| **Eng598/22** | - | - | - | - | - | A | I | - | - | - | - | - | - | - | - | - | - | - |
| **Che854/18** | - | - | - | N | - | - | - | **N** | - | - | - | Q | - | - | - | V | V | V |
| **Agha/17** | - | - | K | - | L | M | - | **N** | E | - | N | Q | N | I | K | V | - | V |

